# Tumor-Cell Invasion Initiates at Invasion Hotspots, an Epithelial Tissue-Intrinsic Microenvironment

**DOI:** 10.1101/2021.09.28.462102

**Authors:** Rei Kobayashi, Hiroaki Takishima, Sheng Deng, Yasuyuki Fujita, Yoichiro Tamori

## Abstract

Malignant cancers emerge in epithelial tissues through a progressive process in which a single transformed mutant cell becomes tumorigenic and invasive. Although numerous genes involved in the malignant transformation of cancer cells have been described, how tumor cells launch an invasion into the basal side of epithelial tissues remains elusive. Here, using a *Drosophila* wing imaginal disc epithelia, we show that genetically mosaic clones of cells mutant for a neoplastic-tumor-suppressor gene (*nTSG*) in combination with the oncogenic Ras (*Ras^V12^*) expression initiate invasion into the basal side of the epithelial layer at specific spots in the epithelial tissue. In this “invasion hotspot”, the oncogenic double-mutant cells activate c-Jun N-terminal kinase (JNK) signaling, which causes basal extrusion of the double-mutant cells and destruction of basement membrane through upregulation of a matrix metalloprotease, MMP1. Conversely, in other regions of the epithelial tissue, the double-mutant cells do not strongly activate JNK, deviate from the apical side of the epithelial layer, and show benign tumor growth in the lumen. These data indicate that the onset of tumor-cell invasion is highly dependent on the tissue-intrinsic local microenvironment. Given the conservation of genetic signaling pathways involved in this process, initiation of tumor-cell invasion from invasion hotspots in *Drosophila* wing imaginal epithelia could help us to understand the developmental mechanisms of invasive cancers.

## 1. Introduction

An epithelial tumor generally originates from a single transformed mutant cell among the highly organized layer of cells which compose the epithelial tissue[1]. If the genetic mutation causes activation of an oncogene or inactivation of a tumor-suppressor gene, the mutant cell will become a pro-tumor cell with the potential to be cancerous. Such nascent pro-tumor cells that emerged within an epithelial layer would evolve into malignant cells with metastatic phenotypes through subsequent transformations over time[1–4].

Tumor development entails a progressive disruption of tissue organization and unleashed proliferation. This indicates that tumor cells deteriorate tissue integrity or evade the robustly organized tissue environment in tumorigenesis[3]. Despite the deterioration of tissue structures, if a tumor grows at the local place and does not spread to other tissues, the tumor can be considered benign. In other words, metastasis from the primary site is the crucial event in cancer progression that transforms a locally growing benign tumor into malignant neoplasms and a life-threatening disease[1].

The first step of the metastatic cascade is invasion, in which tumor cells leave the epithelial layers, penetrate the underlying basement membrane, and migrate through the extracellular matrix (ECM) into the surrounding tissue[5, 6]. The tumor-cell invasion includes various cellular activities such as activation of signaling pathways that control cytoskeletal dynamics and promote cellular survival, turnover of cell-cell and cell-matrix junctions, epithelial-mesenchymal transition (EMT), and proteolysis-dependent ECM degradation, followed by active tumor cell migration into the adjacent tissue[7, 8]. Although numerous genes and signaling pathways involving these different aspects in tumor-cell invasion have been identified, how certain mutant cells escape from the epithelial layer and what cellular and molecular events occur to launch invasive behaviors *in vivo* tissues remain largely elusive[4, 7].

Recent studies especially using the genetically mosaic analysis tools in *Drosophila melanogaster* have greatly contributed to better understanding of the molecular and cellular mechanisms of the early cancer development *in vivo*[9–11]. For example, genetic experiments in *Drosophila* have revealed that the emergence of transformed pro-tumor cells within a normal epithelial layer leads to complex interactions between pro-tumor cells and healthy neighbors[12]. One of such interactions is cell competition, a competitive cellular interaction which occurs when neighboring cells differ in intrinsic cellular properties contributing to selective elimination of either cell type[12–14]. Studies in *Drosophila* epithelial tissues such as developing imaginal discs have shown that genetically mosaic clones mutant for a group of tumor-suppressor genes identified in *Drosophila* – *lethal giant larvae* (*lgl*), *discs large* (*dlg*), and *scribble* (*scrib*)) – are outcompeted by normal neighbors and are therefore eliminated from host tissues[15–18]. Similar cell competition-dependent cell death and elimination of Scribble-knockdown cells have also been demonstrated in mammalian cells using the Madin-Darby Canine Kidney (MDCK) epithelial cell line[19, 20]. These tumor suppressor genes play key roles in the formation of apicobasal cell polarity and regulation of the planar spindle orientation during mitotic cell division in developing epithelial tissues like *Drosophila* imaginal discs[21–24]. When imaginal-disc epithelial cells in *Drosophila* larvae have a homozygous mutation for any of these three genes, the normally monolayered epithelium loses its organized structure, fails to differentiate, and overproliferates thus becoming a multilayered amorphous mass that fuses with adjacent tissues[22]. Loss or alteration in expression of the homologs of these genes in mammals including humans is also associated with tumor development[25–28]. The neoplastic phenotypes exhibited by mutant tissues led to the classification of these three genes as conserved neoplastic tumor-suppressor genes (nTSGs)[22].

The fact that pro-tumor cells like *nTSG* mutants are eliminated by cell competition will closely relate to the data showing cancers arise through the sequential accumulation of multiple mutations in oncogenes and tumor suppressor genes[29, 30]. Indeed, ectopic activation of oncogenic signaling pathways or genes such as *Notch*, JAK/STAT (Janus kinase/signal transducer and activator of transcription), or *Yorkie* (*Yki*: *Drosophila* homolog of Yes-associated protein, YAP) in *nTSG* mutant cells cooperatively induces tumorigenesis[15,18,31–33]. Among these oncogenes, an activated mutant form of Ras small GTPase, *Ras^V12^*, in combination with *nTSG* mutant cells causes highly invasive tumor phenotypes in *Drosophila* and mice[15,26,32,34,35].

In this study, however, we show through detailed analyses of tumor cell phenotypes in *Drosophila* imaginal epithelia that the double mutant cells with a combinatorial mutation of *nTSG* and the oncogenic *Ras* frequently induce apical outgrowth and develop into benign tumors in the lumen. At the same time, the double-mutant cells induce basal extrusion and invasive behaviors only at a few specific spots in the wing imaginal epithelia. These data suggest that the benign-or-malignant fate of tumor cells is highly dependent on a tissue-intrinsic microenvironment and show how the tumor cells begin invasion *in vivo* epithelial tissues.

## 2. Results

### 2.1. nTSG-Ras^V12^ Double Mutant Cells Show Two Morphologically Distinct Tumor Phenotypes

It has been shown that a combinatorial mutation of a neoplastic-tumor-suppressor gene (*nTSG*) and the oncogenic Ras (*Ras^V12^*) induces malignant tumor phenotypes such as intense proliferation and metastatic behaviors in *Drosophila* epithelial tissues[31, 32]. For example, genetically mosaic mutant clones of *scribble* (*scrib*, one of the *nTSG*s) expressing *Ras^V12^* (*scrib-Ras^V12^* clones) generated in the eye imaginal discs and the optic lobes of *Drosophila* larvae show tumorigenic overgrowth and invade the ventral nerve cord[15, 32]. To confirm that the genetically mosaic *scrib-Ras^V12^* clones show the invasive tumor phenotypes in *Drosophila* wing imaginal discs as previously reported in eye imaginal discs[15], we generated the double mutant *scrib-Ras^V12^* clones in the wing imaginal discs using mosaic analysis with a repressible cell marker (MARCM) system[36]. Interestingly, we found that the *scrib-Ras^V12^* clones showed two morphologically distinct tumor phenotypes in the wing imaginal discs. The *scrib-Ras^V12^* clones showed tumor growth at the apical side of the hinge region, so-called “tumor hotspots”[33], but they were in a rounded spherical configuration without protrusions (Figure 1a, c−f). On the other hand, those clones localized at the basal side of the epithelial layer intensely projected cellular protrusions and formed irregularly stretched amorphous shapes (Figure 1b−f).

**Figure 1.**
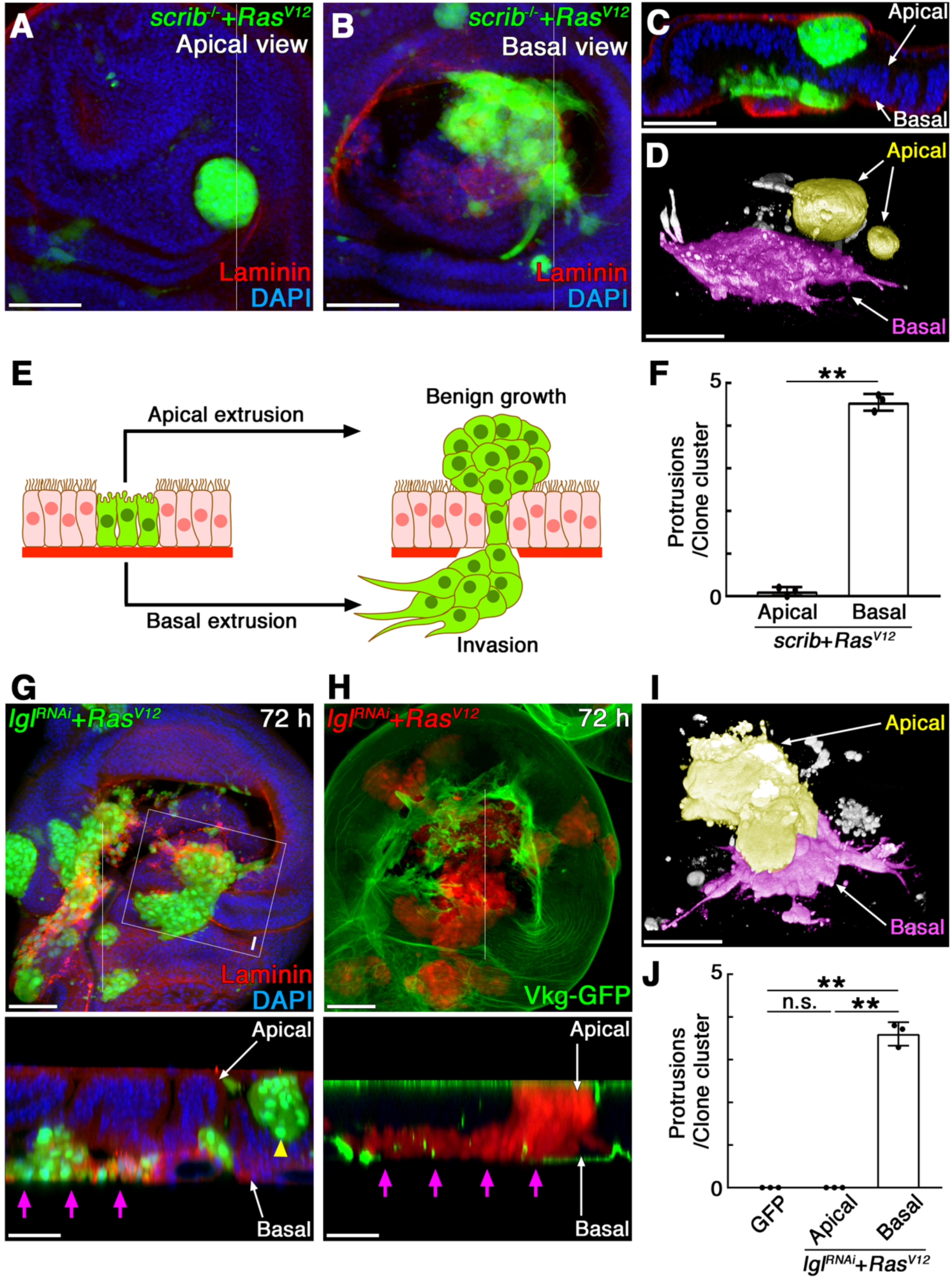
*nTSG-Ras^V12^* double mutant cells show two morphologically distinct tumor phenotypes in wing imaginal epithelia. (**A***−***B**) A wing imaginal disc with *scrib* mosaic mutant clones expressing *Ras^V12^* (labeled by GFP expression: green) four days after clone induction, stained for Laminin-*γ* (red). The images are z-stack projections of 30 confocal images of the apical side (**A**) or basal side (**B**) of the columnar epithelial layer. (**C**) A vertical section at a site indicated by a white line in (**A**) and (**B**). (**D**) A three-dimensional confocal image of GFP signal in (**A**) and (**B**) showing the shapes of mutant clones. The mutant clones located at the apical and basal side of the epithelial layer were pseudocolored with yellow and magenta respectively. (**E**) Schematic representation of tumor phenotypes of *nTSG-Ras^V12^* double mutant cells (green) observed in wing imaginal epithelia. Mutant clones and wild-type cells are shown in green and pink respectively. Basement membranes are drawn as a thick red line. (**F**) Quantification of pseudopodial protrusions in *scrib*-*Ras^V12^* mutant clones (the number of protrusions per mutant clone cluster). Data are mean ± s.d. from three independent experiments. **P<0.001 (unpaired two-tailed Student’s *t*-test); (n=18 GFP-expressing clone clusters from 15 wing discs). (**G***−***H**) Wing discs with mosaic mutant clones coexpressing *lgl^RNAi^* and *Ras^V12^* at the indicated time point after clone induction. Mutant clones were marked with GFP expression (green) in (**G**) and RFP expression (red) in (**H**). Basement membranes were labeled with anti-Laminin *γ* antibody (red) in (**G**) and Vkg-GFP (green) in (**H**). Lower panels: vertical sections at a site indicated by a white line in each upper panel. (**I**) A three-dimensional confocal image of GFP signal showing the shapes of mutant clones indicated by a white square in (**G**). The mutant clones located at the apical and basal side of the epithelial layer were pseudocolored with yellow and magenta respectively. (**J**) Quantification of pseudopodial protrusions in *lgl^RNAi^*-*Ras^V12^* mutant clones (the number of protrusions per mutant clone cluster). Data are mean ± s.d. from three independent experiments. **P<0.001 (unpaired two-tailed Student’s *t*-test); (n=19 GFP-expressing clone clusters from 15 wing discs for each genotype). Nuclei were labeled with DAPI (blue) in (**A***−***C**) and (**G**). Scale bars represent 50 μm. A yellow arrowhead: apically extruded mutant clones. Magenta arrows: basally extruded mutant clones.

To analyze the tumor development induced by *nTSG-Ras^V12^* double mutant clones, we also generated the mosaic clones expressing RNAi for *lethal giant larvae* (*lgl*), another nTSGs, in combination with *Ras^V12^* expression (*lgl^RNAi^-Ras^V12^*) in the wing imaginal epithelia using the heat-shock-induced flip-out Gal4 system[37]. Two days after clone induction, we observed that a subset of the *lgl^RNAi^-Ras^V12^* clones was localized at the apical or basal side of the epithelial layers (Figure S1a*−*c). Three days after clone induction, the *lgl^RNAi^-Ras^V12^* clones in the wing imaginal discs showed two morphologically distinct tumor phenotypes as we observed in the *scrib-Ras^V12^* clones (Figure 1a, b). While the *lgl^RNAi^-Ras^V12^* clones localized at the apical side of tumor hotspots formed round-shape benign-looking tumors without pseudopodial protrusions, those clones localized at the basal side of the epithelial layer formed stretched amorphous shapes and projected pseudopodial protrusions (Figure. 1g, i, j). The basement membrane labeled with anti-laminin *γ* antibody or collagen IV-GFP (Vkg-GFP) was degraded under these basally extruded clones with pseudopodial protrusions (Figure. 1g, h). Four days after clone induction, the *lgl^RNAi^-Ras^V12^* clones grew larger, but the phenotypic differences between the apical and basal tumor clones were clearly observed (data not shown). In the same experimental condition, the mosaic clones expressing only GFP did not show any tumor phenotypes (Figure S1d*−*f). These observations suggest that the *nTSG-Ras^V12^* double mutant clones deviated from the apical side of the epithelial layer develop into benign tumors in the lumen, but they present with invasive behaviors when they are extruded from the basal side (Figure 1e).

When the mosaic clones of *nTSG*-deficient cells without *Ras^V12^* are extruded from the basal side of epithelial layers, they undergo apoptosis and do not show tumorigenic phenotypes[23, 33]. Therefore, we inferred that *Ras^V12^* expression is the primary cause of the invasive phenotypes shown by the *nTSG-Ras^V12^* clones. The phenotypes of *Ras^V12^*-expressing mosaic clones without an *nTSG* defect (*Ras^V12^* clone), however, were different from those observed in the *nTSG-Ras^V12^* clones. Two days after clone induction, a subset of the *Ras^V12^* clones in the wing pouch area was localized at the basal side of the epithelial layer and not labeled by the anti-cleaved DCP1 (Death caspase-1, an effector caspase of *Drosophila*) antibody (Figure. 2a, d). These basally extruded *Ras^V12^* clones forming a cyst-like structure in a spherical shape were localized between the epithelial layer and the underlying basement membrane and did not frequently show pseudopodial protrusions (Figure. 2a, f). Some clones located in the hinge area were localized at the apical side of the epithelial layer and did not show apoptosis (Figure. 2a, d). Three days after clone induction, the basally extruded *Ras^V12^* clones grew larger, did not show apoptosis, but remained to be in a cyst-like structure, and stay between the epithelial layer and the basement membrane (Figure. 2b, e). Although we found a few *Ras^V12^* clones located in the hinge regions showed psudopodial protrusions at the basal side (Figure 6a), these observations suggest that *Ras^V12^* clones, in most settings, undergo benign tumor growth at the basal side of the epithelial layer. Similarly, the apically extruded *Ras^V12^* clones in the hinge area were growing without pseudopodial protrusions at the lumen (Figure. 2b, e).

**Figure 2.**
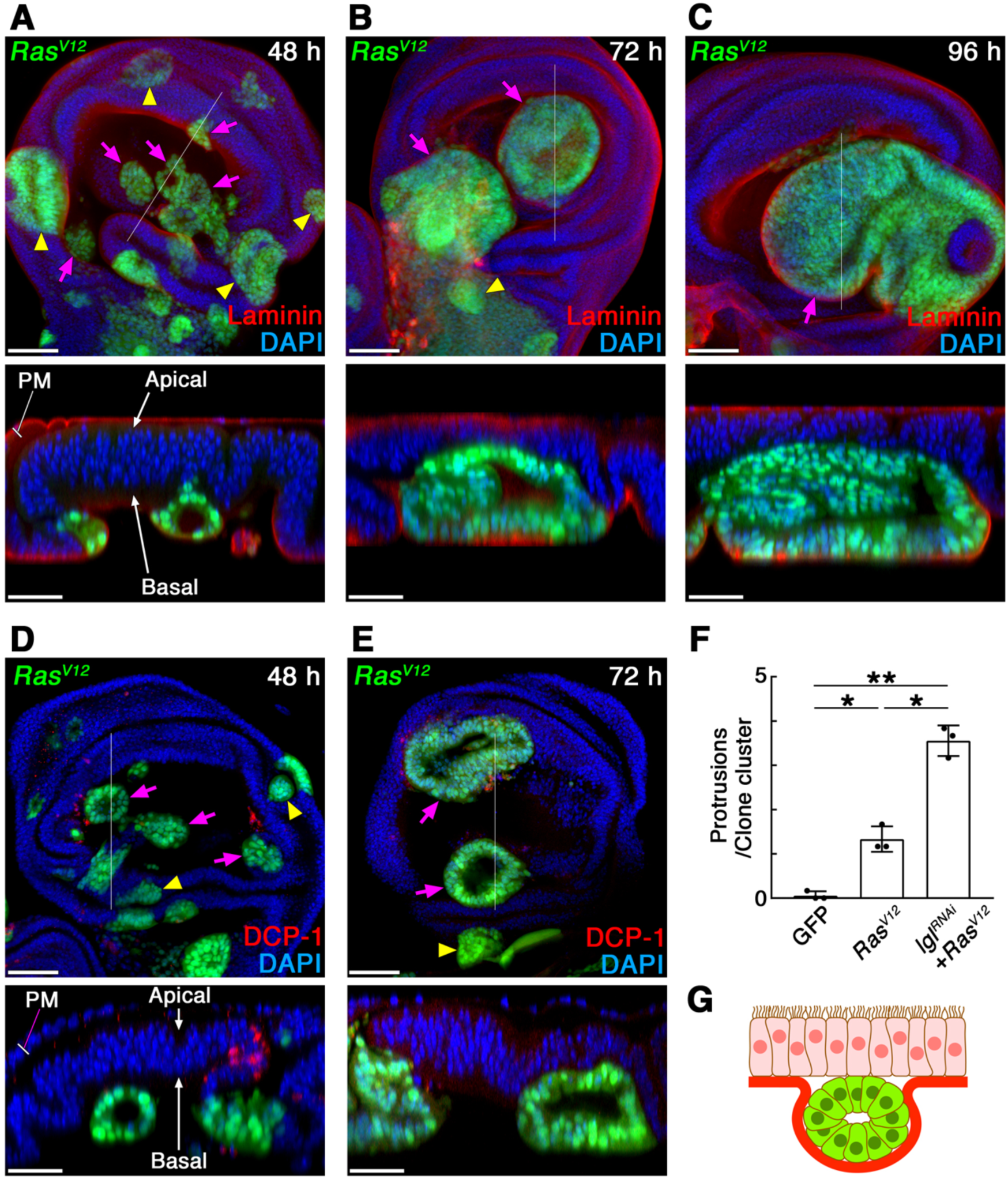
*Ras^V12^*-expressing cells grow as benign tumors in wing imaginal epithelia. (**A***−***C**) Wing imaginal discs with mosaic mutant clones expressing *Ras^V12^* (labeled by GFP expression: green) at the indicated time point after clone induction, stained for Laminin-*γ* (red). The images are z-stack projections of 30 confocal images of the columnar epithelial layer. Lower panels: vertical sections at a site indicated by a white line in each upper panel. (**D***−***E**) Wing imaginal discs with mosaic mutant clones expressing *Ras^V12^* (labeled by GFP expression: green) at the indicated time point after clone induction, stained for cleaved DCP-1 (red). The images are z-stack projections of 30 confocal images of the columnar epithelial layer. Lower panels: vertical sections at a site indicated by a white line in each upper panel. Nuclei were labeled with DAPI (blue) in (**A***−***E**). Scale bars represent 50 μm. Yellow arrowheads: apically extruded mutant clones. Magenta arrows: basally extruded mutant clones. PM: peripodial membrane. (**F**) Quantification of pseudopodial protrusions in the basally extruded mutant clones (the number of protrusions per mutant clone cluster). Data are mean ± s.d. from three independent experiments. *P<0.005, **P<0.001 (unpaired two-tailed Student’s *t*-test); (n=18 GFP-expressing clone clusters from 15 wing discs for each genotype). (**G**) A schematic showing the phenotype of *Ras^V12^*-expressing cells in wing imaginal epithelia. Mutant clones and wild-type cells are shown in green and pink respectively. Basement membranes are drawn as a thick red line.

### 2.2. JNK-MMP1 Signaling Is Activated in the Basally Invading nTSG-Ras^V12^ Tumor Clones

The morphological observations of the *nTSG-Ras^V12^* mosaic clones lead us to reason that those two distinct tumor phenotypes shown by the double-mutant cells are dependent on the location in the epithelial tissue; a benign tumor growth at the apical side, and a malignant invasive phenotype at the basal side of the epithelial layer (Figure 1e). One of the distinctive signs of invasive tumor cells in epithelial tissues is upregulated expression of matrix metalloproteinases (MMPs), endopeptidases which are capable of degrading ECM proteins[38]. *Drosophila* has two MMPs (MMP1 and MMP2), and MMP1 has been shown to be involved in the invasive phenotypes of tumor cells[39]. Immunofluorescence using anti-*Drosophila* MMP1 antibody revealed that the MMP1 expression was strongly upregulated in the *lgl^RNAi^-Ras^V12^* clones localized at the basal side of the epithelial layer (Figure. 3a, b, g). By contrast, the spherical tumor clones growing at the apical side of the epithelial layer did not show strong upregulation of MMP1 expression (Figure. 3a, b, g). In the same experimental condition, the mosaic clones expressing only GFP did not show upregulation of MMP1 expression (Figure S2a, c).

**Figure 3.**
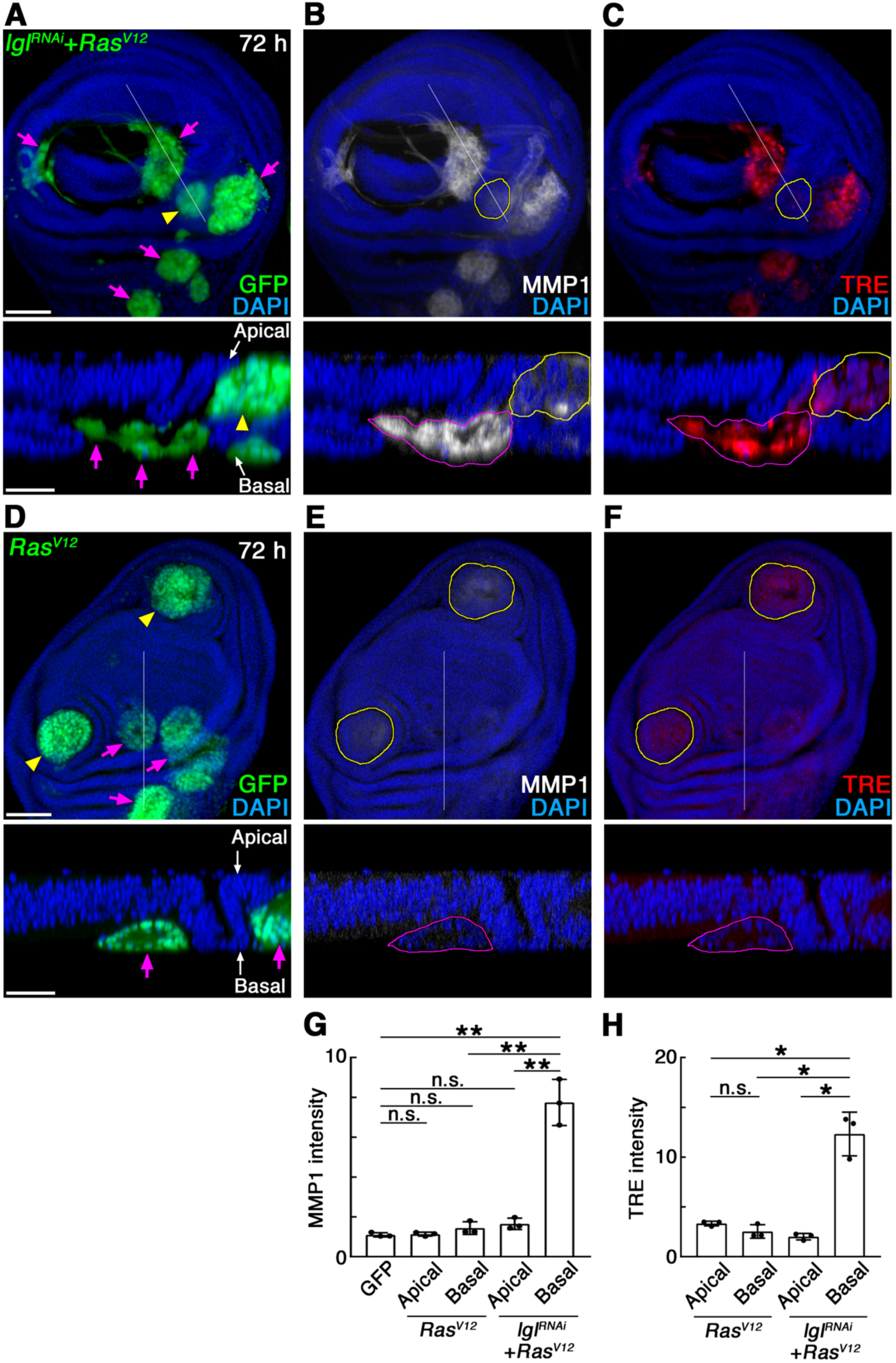
JNK-MMP1 signaling is activated in the basally invading *lgl^RNAi^-Ras^V12^* tumor clones. (**A***−***C**) A wing imaginal disc with mosaic mutant clones expressing *lgl^RNAi^* and *Ras^V12^* 72 hours after clone induction. The mutant clones are labeled by GFP expression (green) in (**A**). (**B**) MMP1 expression detected by anti-MMP1 antibody staining (white) in the mosaic wing disc in (**A**). (**C**) JNK activation detected by TRE-DsRed in the mosaic wing disc in (**A**). The images are z-stack projections of 30 confocal images of the columnar epithelial layer. Lower panels: vertical sections at a site indicated by a white line in each upper panel. (**D***−***F**) Wing imaginal discs with mosaic mutant clones expressing *Ras^V12^* 72 hours after clone induction. The mutant clones are labeled by GFP expression (green) in (**D**). (**E**) MMP1 expression detected by anti-MMP1 antibody staining (white) in the mosaic wing disc in (**D**). (**F**) JNK activation detected by TRE-DsRed in the mosaic wing disc in (**D**). Apically or basally extruded clones are circled by yellow or magenta lines respectively in (**B**, **C**, **E**, **F**). The images are z-stack projections of 30 confocal images of the columnar epithelial layer. Lower panels: vertical sections at a site indicated by a white line in each upper panel. Nuclei were labeled with DAPI (blue) in (**A***−***F**). Scale bars represent 50 μm. Yellow arrowheads: apically extruded mutant clones. Magenta arrows: basally extruded mutant clones. (**G***−***H**) Quantification for signal intensities of anti-MMP1 antibody staining (**G**) or TRE-DsRed (**H**). Values are expressed as a ratio relative to control (GFP-negative areas). Data are mean ± s.d. from three independent experiments. *P<0.005, **P<0.001 (unpaired two-tailed Student’s *t*-test); n=90 selected areas of 25-square pixels (for MMP1) or 225-square pixels (for TRE-DsRed) from GFP-positive clone regions.

In the context of tumor progression, MMP1 expression is induced by activation of the c-Jun N-terminal kinase (JNK) signaling pathway in *Drosophila* epithelial tissues[39]. Thus, we examined the activity of the JNK signaling pathway in the *nTSG-Ras^V12^* double-mutant clones using TRE-DsRed, a JNK signaling reporter[40]. The TRE-DsRed signals showed us that the JNK activation pattern was exactly similar to the MMP1 expression pattern in the mosaic mutant clones. The signal of TRE-DsRed was strongly upregulated in the *lgl^RNAi^-Ras^V12^* clones localized at the basal side of the epithelial layer, but the signal was weak in these clones localized at the apical side (Figure. 2a, c, h). In the same experimental condition, the mosaic clones expressing only GFP did not show upregulation of the TRE-DsRed signal (Figure S2a, b). These results suggest that JNK-MMP1 signaling are strongly activated when the double-mutant cells are extruded from the basal side of epithelial layers.

### 2.3. JNK-MMP1 Signaling Is Activated by the nTSG Defect

The morphological observations of *Ras^V12^*-expressing mosaic clones without an *nTSG* defect showed that these clones do not induce invasive phenotypes in most cases even at the basal side of the epithelial layer (Figure. 2). To corroborate this, we examined the expression patterns of MMP1 and TRE-DsRed in the wing imaginal discs with *Ras^V12^* mosaic clones. Three days after clone induction, neither MMP1 expression nor TRE-DsRed signals were observed in the *Ras^V12^* clones growing at the apical side of the epithelial layer (Figure 3d−f). Although a few *Ras^V12^* clones located at the basal side of the epithelial layer infrequently showed expressions of MMP1 and TRE-DsRed, most of the clones did not show strong upregulation of these even at the basal side of the epithelial layers (Figure 3e−h). These results suggest that misexpression of *Ras^V12^* itself does not induce JNK-MMP1 signaling activation in the epithelial tissues.

Then, what does cause JNK activation in the basally invading *nTSG-Ras^V12^* double-mutant cells? It has been reported that Grindelwald, a *Drosophila* tumor necrosis factor (TNF) receptor, integrates signals from both TNF and apical polarity determinants to induce JNK activation in response to perturbation of epithelial apicobasal polarity[42]. Thus, we asked whether the JNK-MMP1 signaling activation observed in the *nTSG-Ras^V12^* double mutant tumor cells is caused by the *nTSG* defect-induced apicobasal polarity disruption. To address this question, we tested JNK-MMP1 signaling activities in the *lgl^RNAi^-*mosaic clones without *Ras^V12^* expression. It has been previously reported that genetically mosaic clones of *nTSG*-deficient cells, such as *lgl*- or *scrib*-mutant clones, show apoptosis as the result of cell competition when they are surrounded by normal cells in epithelial tissues[17]. In the process of cell competition-induced apoptosis, JNK signaling plays a key role to activate the caspase-signaling pathway in loser cells[15]. When the *nTSG*-deficient cells are not surrounded by normal wild-type cells, they can survive, disrupt epithelial tissue organization, and show tumorigenic overgrowth. JNK signaling has also been shown to be involved in the process of apicobasal polarity defect-induced tumorigenesis[42].

We have previously shown that *lgl^RNAi^-*mosaic clones without *Ras^V12^* expression showed apoptosis and were extruded toward the basal side of the epithelial layer[33]. To analyze the tumor phenotypes induced by an *nTSG* defect and the JNK signaling activity in its process, we kept the *lgl^RNAi^-*mosaic clones alive by blocking their apoptosis with a co-expression of *p35*, an anti-apoptotic gene of baculovirus. Three days after mosaic clone induction, a subset of the clones co-expressing *lgl^RNAi^* and *p35* (*lgl^RNAi^*-*p35* clones) were localized at the basal side of the epithelial layer and show mild proliferation (Figure. 4a). Although the basement membrane surrounding these *lgl^RNAi^*-*p35* clones was partially degraded, these clones did not show pseudopodial protrusions (Figure. 4a, b). Four days after clone induction, most of the *lgl^RNAi^*-*p35* clones were localized at the basal side of the epithelial layer, and we found that both JNK activity and MMP1 expression were upregulated in these basally extruded clones (Figure 4e−l). These basally extruded clones, however, did not frequently show pseudopodial protrusions (Figure. 4b, g, k). Although we found that the apically extruded *lgl^RNAi^*-*p35* clones showed weak expression of TRE-DsRed, MMP1 expression was hardly observed in these apical clones (Figure 4c, h, j). These results suggest that activation of the JNK-MMP1 signaling is induced by an *nTSG* defect but is not enough to induce the invasion phenotypes such as pseudopodial protrusions observed in the *nTSG-Ras^V12^* double mutant cells (Figure 4d).

**Figure 4.**
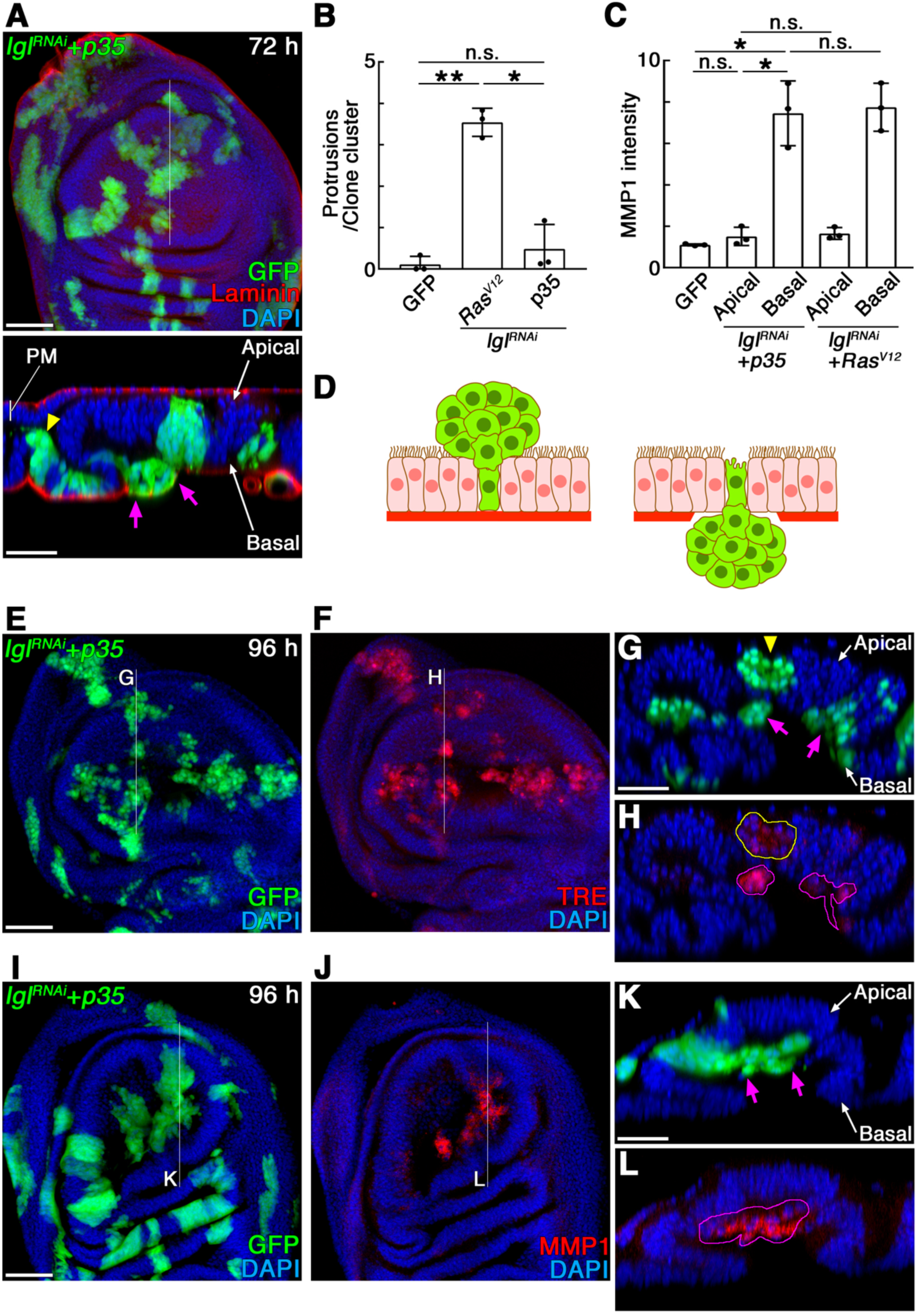
*lgl^RNAi^-p35* clones induce basal extrusion and JNK-MMP1 activation but not invasive behaviors. (**A**) A wing imaginal disc with mosaic mutant clones expressing *lgl^RNAi^* and *p35* (labeled by GFP expression: green) 72 hours after clone induction, stained for Laminin-*γ* (red). The images are z-stack projections of 30 confocal images of the columnar epithelial layer. Lower panel: a vertical section at a site indicated by a white line in the upper panel. PM: peripodial membrane. (**B**) Quantification of pseudopodial protrusions in the basally extruded mutant clones (the number of protrusions per mutant clone cluster). Data are mean ± s.d. from three independent experiments. *P<0.005, **P<0.001 (unpaired two-tailed Student’s *t*-test); (n=20 GFP-expressing clone clusters from 15 wing discs for each genotype). (**C**) Quantification for signal intensities of anti-MMP1 antibody staining. Values are expressed as a ratio relative to control (GFP-negative areas). Data are mean ± s.d. from three independent experiments. *P<0.005 (unpaired two-tailed Student’s *t*-test); n=90 selected areas of 25-square pixels from GFP-positive clone regions. (**D**) Schematics showing the phenotypes of *lgl^RNAi^*-*p35* clones in wing imaginal epithelia. Left: apically extruded clones. Right: basally extruded clones. Mutant clones and wild-type cells are shown in green and pink respectively. Basement membranes are drawn as a thick red line. (**E**) Wing imaginal discs with mosaic mutant clones expressing *lgl^RNAi^* and *p35* 96 hours after clone induction. The mutant clones were labeled by GFP expression (green). (**F**) JNK activation detected by TRE-DsRed in the mosaic wing disc in (**E**). (**G***−***H**) A vertical section at a site indicated by a white line in (**E***−***F**). (**I**) Wing imaginal discs with mosaic mutant clones expressing *lgl^RNAi^* and *p35* 96 hours after clone induction. The mutant clones were labeled by GFP expression (green). (**J**) MMP1 expression detected by anti-MMP1 antibody staining (red) in the mosaic wing disc in (**I**). The images are z-stack projections of 30 confocal images of the columnar epithelial layer. (**K***−***L**) A vertical section at a site indicated by a white line in (**I***−***J**). Nuclei were labeled with DAPI (blue) in (**A**) and (**E***−***L**). A yellow arrowhead and circle: apically extruded mutant clones. Magenta arrows and circles: basally extruded mutant clones. Scale bars represent 50 μm.

### 2.4. JNK Activation Is Involved in the Extrusion of nTSG-Deficient Cells

To gain further insight into the role of the JNK signaling pathway in the invasive phenotype of *nTSG-Ras^V12^* mutant cells, we suppressed JNK activity in the *lgl^RNAi^-Ras^V12^* double mutant clones. When a dominant-negative form of *basket* (*Drosophila* JNK), *bsk^DN^*, was co-expressed in the *lgl^RNAi^-Ras^V12^* mosaic clones, they proliferated in the epithelial layer, induced curvature of the epithelial layer, but did not show extrusions even three days after clone induction (Figure 5a). We also confirmed that MMP1 expression was suppressed in these clones (Figure 5b). When we suppressed the function of MMP1 in the *lgl^RNAi^-Ras^V12^* mosaic clones by co-expression of *Drosophila* Tissue inhibitor of metalloproteinases, Timp, however, *lgl^RNAi^-Ras^V12^* clones showed basal extrusion at three days after clone induction (Figure 5c). Although the basement membrane under the growing mutant clones was partially degraded, these clones were localized between the basal side of the epithelial layer and the basement membrane and did not show pseudopodial protrusions (Figure 5c, e, f). This result indicates that MMP1 is not involved in the basal extrusion. Based on these data, we reasoned that JNK activation promotes two independent downstream events: basal extrusion of *nTSG-Ras^V12^* clones[43] and MMP1 expression-induced basement membrane degradation[39]. The mechanism of the JNK-dependent extrusion of *nTSG* mutant cells has been shown in *Drosophila* eye imaginal discs; the signaling of Slit ligand, its transmembrane Roundabout receptor Robo2, and the downstream cytoskeletal effector Enabled/VASP (Ena) exert a force downstream of JNK to induce delamination of *scrib* mutant cells from epithelial layers[43]. As we demonstrated above, however, the *Ras^V12^*-expressing clones without an *nTSG* defect underwent basal extrusion as a cell cluster without JNK activation (Figure. 3f). Therefore, the cell-cluster extrusion of *Ras^V12^* clones is not dependent on the JNK-signaling activity but might be induced by defective epithelial morphogenesis[41].

**Figure 5.**
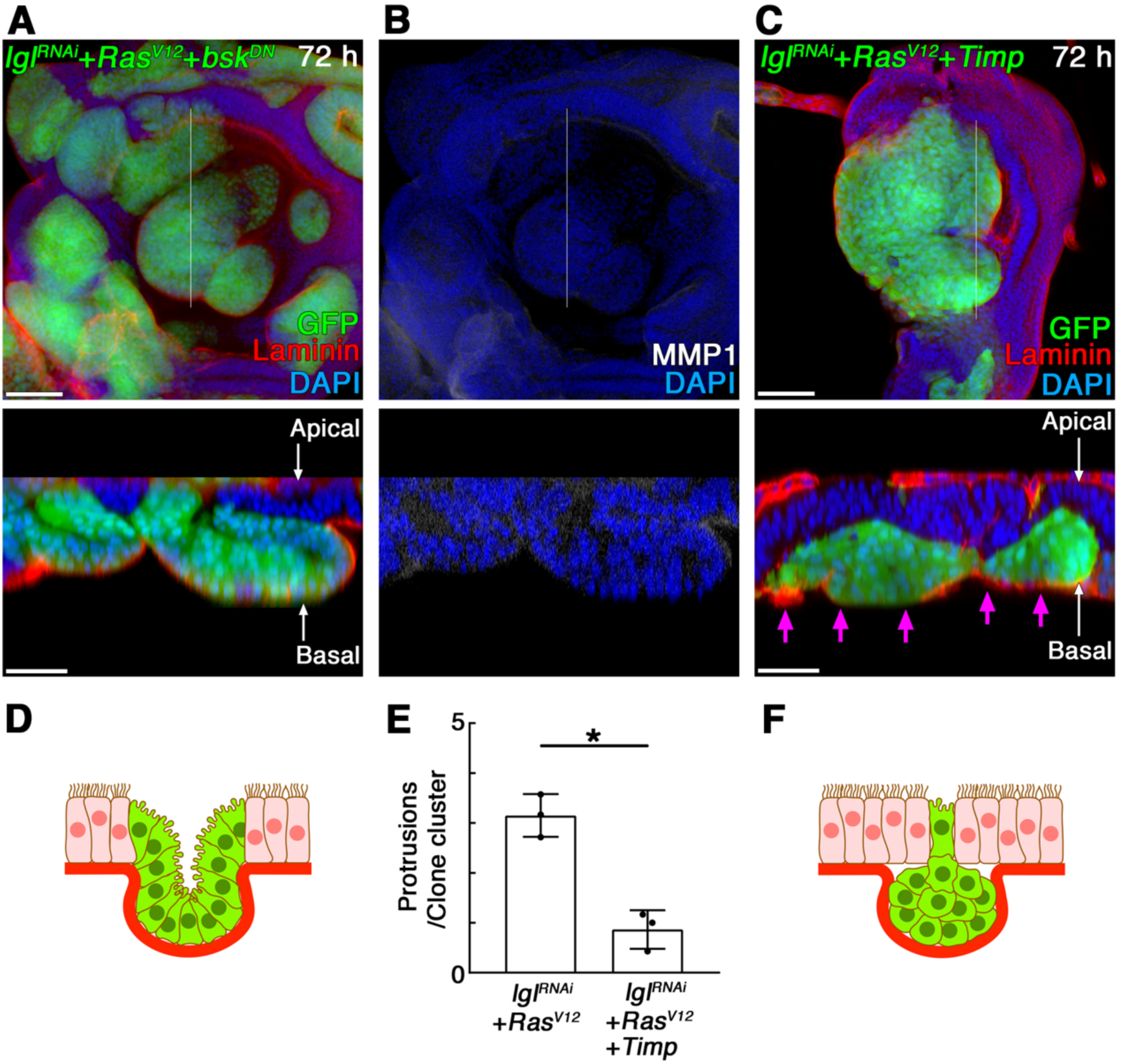
JNK activation but not MMP1 is involved in the basal extrusion of *lgl^RNAi^*-*Ras^V12^* clones. (**A**) A wing imaginal disc with mosaic mutant clones expressing *lgl^RNAi^*, *Ras^V12^*, and *bsk^DN^* (labeled by GFP expression: green) 72 hours after clone induction, stained for Laminin-*γ* (red). (**B**) MMP1 expression detected by anti-MMP1 antibody staining (white) in the mosaic wing disc in (**A**). (**C**) A wing imaginal disc with mosaic mutant clones expressing *lgl^RNAi^*, *Ras^V12^*, and *Timp* (labeled by GFP expression: green) 72 hours after clone induction, stained for Laminin-*γ* (red). The images of the upper panels are z-stack projections of 30 confocal images of the columnar epithelial layer. Lower panel: a vertical section at a site indicated by a white line in each upper panel. Magenta arrows indicate basally translocated mutant clones. Nuclei were labeled with DAPI (blue) in (**A***−***C**). Scale bars represent 50 μm. (**D**) A schematic showing the phenotypes of *lgl^RNAi^*-*Ras^V12^*-*bsk^DN^* clones in wing imaginal epithelia. Mutant clones and wild-type cells are shown in green and pink respectively. Basement membranes are drawn as a thick red line. (**E**) Quantification of pseudopodial protrusions in the basally extruded mutant clones (the number of protrusions per mutant clone cluster). Data are mean ± s.d. from three independent experiments. *P<0.005 (unpaired two-tailed Student’s *t*-test); (n=20 GFP-expressing clone clusters from 15 wing discs for each genotype). (**F**) A schematic showing the phenotypes of *lgl^RNAi^*-*Ras^V12^*-*Timp* clones in wing imaginal epithelia. Mutant clones and wild-type cells are shown in green and pink respectively. Basement membranes are drawn as a thick red line.

### 2.5. Epithelial Cell Polarity is Intrinsically Compromised at The Invasion Hotspots

As we demonstrated above, the *lgl^RNAi^*-*p35* clones without *Ras^V12^* expression do not show invasive phenotypes, whereas they activate the JNK-MMP1 signaling and are basally extruded (Figure. 4). Also, the *lgl^RNAi^-Ras^V12^* clones expressing Timp basally translocate, stay between the basal side of the epithelial layer and the basement membrane, but do not show invasive phenotypes (Figure. 5). Similarly, most of the *Ras^V12^*-expressing clones without an *nTSG* defect neither activate JNK nor show invasive phenotypes even at the basal side of epithelial layers (Figure 3). Taking all these data together we hypothesized that both *Ras^V12^* expression and JNK activation are required for the onset of invasive phenotypes and two independent processes lead to the tumor invasion: 1) the JNK activation caused by an apicobasal polarity defect induces basal extrusion of tumor cells and MMP1 upregulation-mediated basement membrane degradation, and 2) oncogenic Ras and the JNK activation cooperatively provoke invasive behaviors of tumor cells at the extracellular matrix.

One of our observations that contradicts this hypothesis is that a few *Ras^V12^*-expressing clones show an upregulation of JNK-MMP1 signaling and invasive phenotypes at the basal side of epithelial layers albeit infrequently (Figure 6a, b). Interestingly, we realized that the infrequent basal invasion phenotypes of *Ras^V12^*-expressing clones were almost always observed at a few specific areas in the wing imaginal epithelia. A quantification for this localized pattern of the basal invasion of *Ras^V12^*-expressing clones revealed that, surprisingly, over 80% of basal invasions were derived from four specific areas located in the hinge region (area number 2, 6, 7, 8 in Figure 6e). Moreover, we found that the basally extruded MMP1-positive *lgl^RNAi^-Ras^V12^* clones were also derived from these specific areas at the almost same ratio (Figure 6c−e). We, therefore, termed these areas “invasion hotspots.” Among these invasion hotspots in the wing imaginal discs, the occurrence ratio of basal invasion phenotypes was relatively high at the presumptive unnamed plate (or humeral plate) area[44] (area number 7 in Figure 6e) in the dorsal hinge region and at the presumptive axillary pouch (or pleural sclerite) area[44] (area number 2 in Figure 6e) in the ventral hinge region.

**Figure 6.**
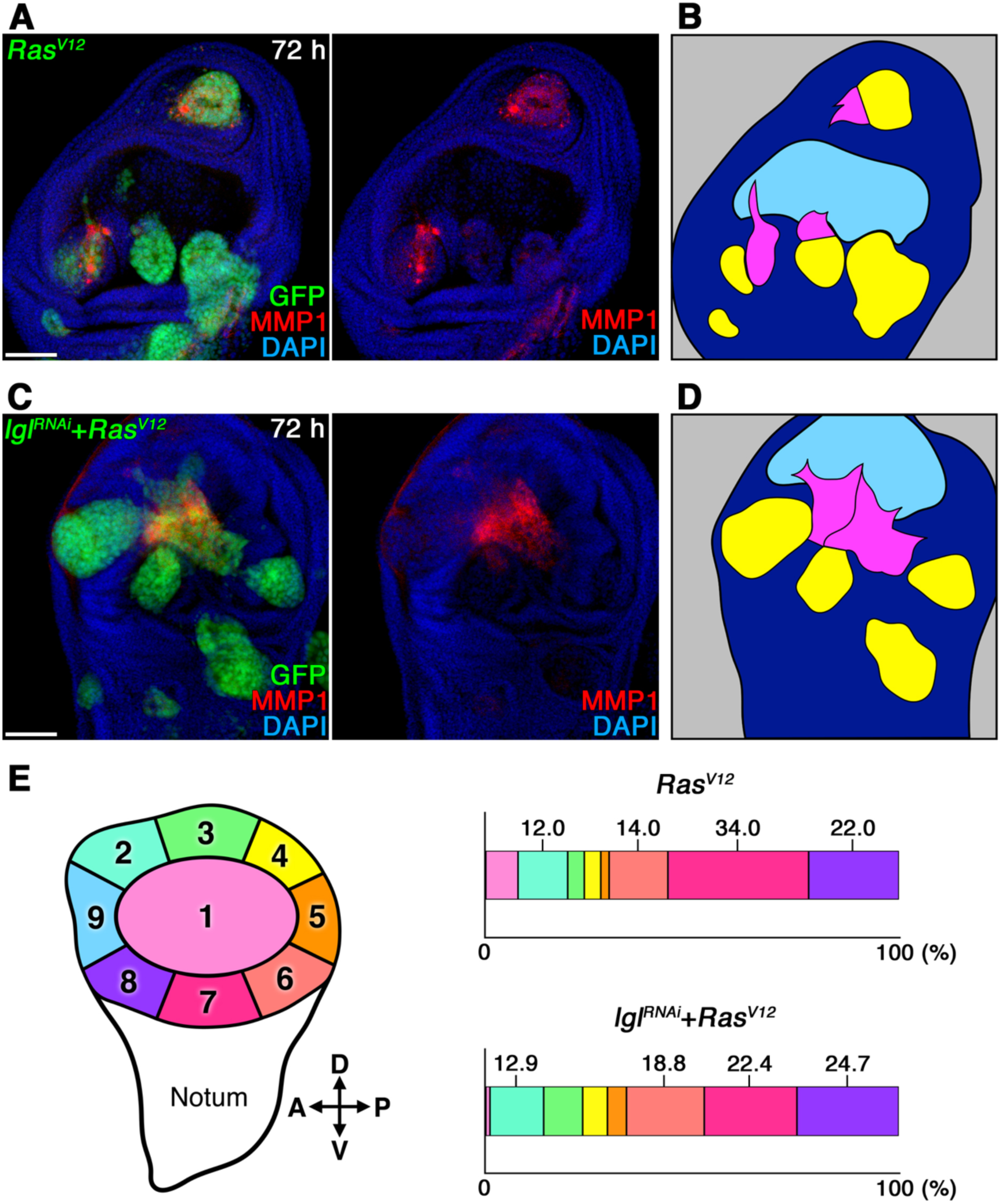
*Ras^V12^*-expressing clones show basal invasion phenotypes at the specific spots in wing disc epithelia. (**A**) A wing imaginal disc with mosaic mutant clones expressing *Ras^V12^* (labeled by GFP expression: green) 72 hours after clone induction, stained for MMP1 (red). The right panel shows the MMP1 expression pattern (red) in the left panel. (**B**) A line drawing traces the tumor clones located at the apical side (yellow) and basal side (magenta) of the wing disc epithelia shown in (**A**). The wing pouch area is shown in light blue. (**C**) A wing imaginal disc with mosaic mutant clones expressing *lgl^RNAi^* and *Ras^V12^* (labeled by GFP expression: green) 72 hours after clone induction, stained for MMP1 (red). The right panel shows the MMP1 expression pattern (red) in the left panel. (**D**) A line drawing traces the tumor clones located at the apical side (yellow) and basal side (magenta) of the wing disc epithelia shown in (**C**). The wing pouch area is shown in light blue. (**E**) Quantification of locational occurrence ratio of basal invasion (basally extruded mutant clones with pseudopodial protrusions) induced by *Ras^V12^*- or *lgl^RNAi^*-*Ras^V12^* mosaic clones in wing disc epithelia. n=50 GFP-expressing clone clusters from 73 wing discs for *Ras^V12^*-clones or n=85 GFP-expressing clone clusters from 54 wing discs for *lgl^RNAi^*-*Ras^V12^* clones.

We presumed that these invasion hotspots have some differences from other regions in intrinsic tissue structures or local genetic signaling activities. One of the apparent differences we found in these spots is a disturbed pattern of planar-polarized cellular arrangement visible from the basal side of the epithelial layer (Figure 7a). This cellular arrangement pattern was visualized by an anti-tubulin antibody staining, and the invasion hotspot was recognized as a disturbance of the flow-like pattern of microtubules (Figure 7a−e). Furthermore, we found that the subcellular localization patterns of adherens junction proteins and cytoskeletal proteins were altered in these invasion hotspots. Immunofluorescences with anti-E-cadherin and anti-Armadillo (*Drosophila β*-catenin) antibodies showed that both proteins were localized not only at the apical side but also at the basal-lateral side of epithelial cells in the invasion hotspots (Figure 7h, j, m, o), whereas the adherens junction normally localized to the apical-lateral membrane (Figure 7g, i, l, n). In addition, the subcellular localization patterns of the actin cytoskeleton (F-actin) and *α*-Spectrin (a subunit of the spectrin cytoskeleton) were shifted from the apical to the basal side in the invasion hotspots (Figure 7p−y). These observations suggest that the epithelial apicobasal polarity is intrinsically compromised in the cells of invasion hotspots albeit mildly and that the spots are susceptible to stimuli which disturb epithelial organization such as oncogenic mutations.

**Figure 7.**
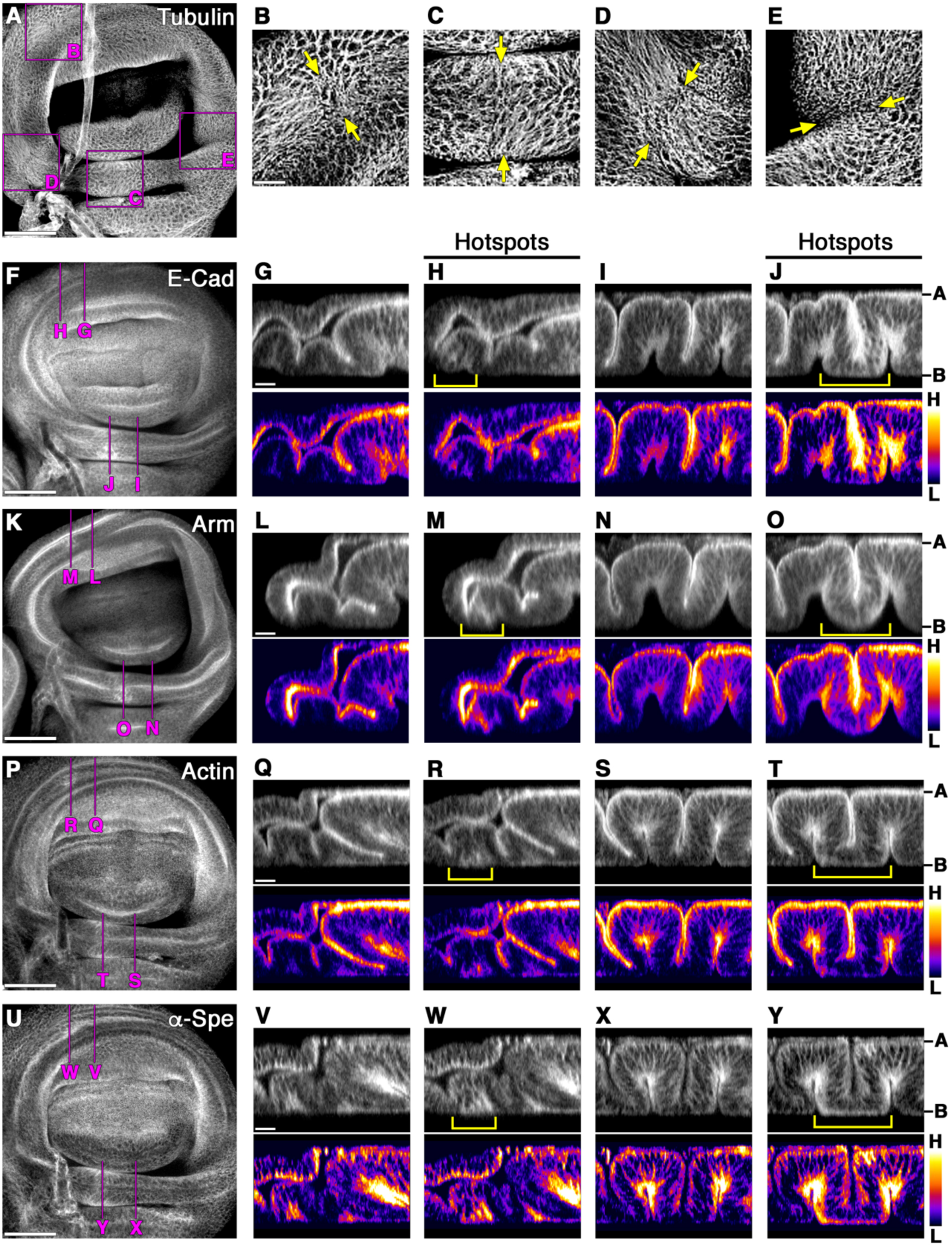
Epithelial cell polarity is intrinsically compromised at the invasion hotspots. (**A**) A wild-type wing disc stained for *α*-tubulin (white). (**B***−***E**) Magnifications of the boxes indicated in (**A**). Yellow arrows indicate the invasion hotspots. (**F**) A wild-type wing disc stained for E-Cadherin (white). (**G***−***J**) Vertical sections at sites indicated by magenta lines in (**F**). Lower panels: Signal intensities of each upper panel image. (**K**) A wild-type wing disc stained for Armadillo (white). (**L***−***O**) Vertical sections at sites indicated by magenta lines in (**K**). Lower panels: Signal intensities of each upper panel image. (**P**) A wild-type wing disc stained for F-actin (white). (**Q***−***T**) Vertical sections at sites indicated by magenta lines in (**P**). Lower panels: Signal intensities of each upper panel image. (**U**) A wild-type wing disc stained for *α*-Spectrin (white). (**V***−***Y**) Vertical sections at sites indicated by magenta lines in (**U**). Lower panels: Signal intensities of each upper panel image. The images in (**A**, **F**, **K**, **P**, **U**) are z-stack projections of 30 confocal images of the basal side of the columnar epithelial layer. Yellow brackets show invasion hotspots in (**H**, **J**, **M**, **O**, **R**, **T**, **W**, **Y**). Scale bars represent 50 μm in (**A**, **F**, **K**, **P**, **U**) and 10 μm in (**B**, **G**, **L**, **Q**, **V**). A: apical. B: basal. H: high. L: low.

### 2.6. Ras^V12^ Expression Induces Basal Invasion Specifically at The Invasion Hotspots

To examine whether *Ras^V12^*-expression induces invasive phenotypes specifically at the invasion hotspots, we used a Gal4-driver line, *GMR17G12-Gal4*, that expresses Gal4 specifically in the entire medial fold of the dorsal hinge area during larval development (Figure 8a−c). When *Ras^V12^*-misexpression was induced by *GMR17G12-Gal4*, basal invasions and pseudopodial protrusions were observed specifically at the invasion hotspots (Figure 8d−f). We also found that the MMP1 expression was specifically upregulated in the invasive cells protruded from these spots (Figure 8g−i). When only RFP expression was induced in the dorsal hinge area with *GMR17G12-Gal4*, neither tumor phenotypes nor MMP1 upregulation was observed (Figure S3a*−*c). These results lead us to conclude that *Ras^V12^*-expression induces basal invasion specifically at the invasion hotspots in the epithelial tissues. When *lgl^RNAi^-Ras^V12^* double mutant was induced in the entire medial fold using *GMR17G12-Gal4*, the JNK-MMP1 activation was observed at the invasion hotspot before the basal invasion occurred (Figure S3d*−*f). Subsequently, *lgl^RNAi^-Ras^V12^* double mutant cells in this area showed basal invasion and intensive pseudopodial protrusions from the invasion hotspots (Figure 8j−l). Collectively, these data suggest that *Ras^V12^*-expression cooperating with the JNK signaling which is prone to be activated at the invasion hotspots provoke invasive phenotypes in the wing imaginal epithelia.

**Figure 8.**
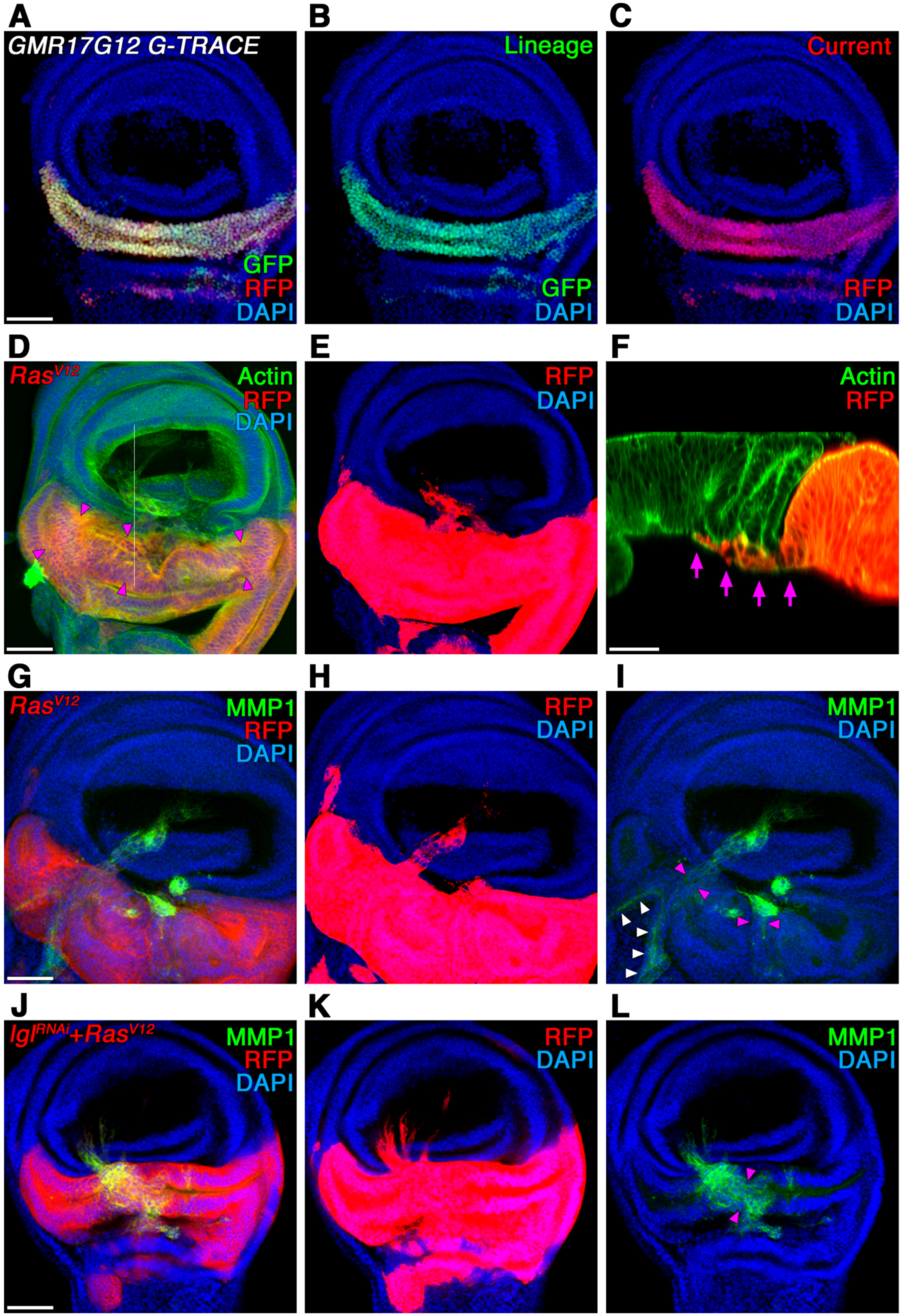
*Ras^V12^* expression induces basal invasion specifically at the invasion hotspots in the wing imaginal epithelia. (**A***−***C**) A wing imaginal disc showing G-TRACE analysis with *GMR17G12-Gal4*. Lineage-traced GFP expression (**B**) and current RFP-expression (**C**) are merged in (**A**). (**D**) A wing imaginal disc with *GMR17G12-Gal4*-driven *Ras^V12^* expression in the dorsal hinge region stained for F-actin (green). *GMR17G12-Gal4* expressing regions were labeled by RFP (red). (**E**) The fluorescent intensity of the RFP signal in (**D**) was increased to visualize pseudopodial protrusions. (**F**) A vertical section at a site indicated by a white line in (**D**). Magenta arrows indicate basal invasion of *Ras^V12^*-expressing cells. (**G**) A wing imaginal disc with *GMR17G12-Gal4*-driven *Ras^V12^* expression in the dorsal hinge region stained for MMP1 (green). *GMR17G12-Gal4* expressing regions were labeled by RFP (red). (**H**) The fluorescent intensity of the RFP signal in (**G**) was increased to visualize pseudopodial protrusions. (**I**) MMP1 expression pattern (green) in the wing disc in (**G**). White arrowheads indicate endogenous MMP1 expression in the trachea. (**J**) A wing imaginal disc with *GMR17G12-Gal4*-driven *lgl^RNAi^* and *Ras^V12^* expression in the dorsal hinge region stained for MMP1 (green). *GMR17G12-Gal4* expressing regions were labeled by RFP (red). (**K**) The fluorescent intensity of the RFP signal in (**J**) was increased to visualize pseudopodial protrusions. (**L**) MMP1 expression pattern (green) in the wing disc in (**J**). Magenta arrowheads indicate invasion hotspots in (**D**), (**I**), and (**L**). Nuclei were labeled with DAPI (blue) in (**A***−***E**) and (**G***−***L**). Scale bars represent 50 μm.

## 3. Discussion

This study describes how a combinatorial mutation of an *nTSG* defect and the oncogenic Ras activation induces tumor invasion *in vivo* using *Drosophila* wing imaginal epithelia as an experimental model. Although the invasive tumor phenotypes of *nTSG-Ras^V12^* double mutant cells have been previously described in different systems[15,26,32,34,35], our data show that their tumor phenotypes, benign tumor growth or malignant invasive phenotypes, are highly dependent on a tissue-intrinsic microenvironment. On the one hand, the *nTSG-Ras^V12^* double mutant cells develop into benign tumors when they are extruded from the apical side of the epithelial layer. On the other, they become invasive when they are extruded from the basal side. Our data suggest that the invasion phenotypes of *nTSG-Ras^V12^* double mutant clones are implemented by the combination of three independent processes: JNK activation-induced basal extrusion, MMP1-mediated basement membrane degradation, and activation of invasive cellular behaviors by oncogenic Ras at the ECM. One of the questions unanswered in this study is why the *lgl^RNAi^-Ras^V12^* double mutant cells do not induce JNK-MMP1 signaling activation and invasive phenotypes at the apical side of the epithelial layer. Based on our data, we can speculate that an environmental factor has a key role in determining the tumor phenotypes. In fact, tumor cell invasion is regarded as an adaptive process mediated by the interactions with stromal components, including ECM, fibroblasts, endothelial cells, and macrophages at the basal side of the epithelial layer[7,45,46]. Conversely, when tumor cells are extruded into the lumen from the apical side of the epithelial layer, they do not have interactions with those stromal components. In *Drosophila* imaginal discs, hemocytes are recruited to the sites where basement membranes are degraded and have an interaction with tumor cells[47–49]. Therefore, tumor cells, after penetrating the basement membrane, encounter a stromal component, supposedly ECM and hemocytes in *Drosophila* imaginal discs, which may further enhance JNK-MMP signaling and provoke invasive behaviors.

We previously reported that *nTSG*-deficient cells (without *Ras^V12^* expression) extruded from the apical side of the epithelium begin tumorigenic overgrowth at the tumor hotspots in wing imaginal discs, whereas those cells extruded toward the basal side at the tumor coldspots undergo apoptosis[33]. It has been well documented that oncogenic Ras has a decremental effect on the apoptosis pathways whereby it contributes to the survival of cancer cells[50, 51]. Mechanistically, oncogenic Ras activates the PI3K-Akt and Raf-MAPK signaling pathways that lead to downregulation of proapoptotic mediators or upregulation of anti-apoptotic genes[51]. These anti-apoptotic functions of oncogenic Ras help the *lgl^RNAi^-Ras^V12^* double mutant cells survive even when they are extruded from the basal side of epithelial layers. Our data also show that the *lgl^RNAi^*-p35 clones (without *Ras^V12^* expression) do not induce invasive phenotypes, whereas they activate the JNK-MMP1 signaling and are basally extruded. Besides the prosurvival signalings such as PI3K-Akt and Raf-MAPK pathways, Ras activates Rho GTPases which play a key role in alterations of cell adhesion and cell motility[51]. Collectively, all these data indicate that the oncogenic Ras-induced alterations of multiple signaling activities are required for the onset of the invasive phenotypes of *nTSG-Ras^V12^* double mutant cells.

In this study, however, we also show that *Ras^V12^*-expressing clones without an *nTSG* defect do not show invasive phenotypes except in the setting of those clones at the invasive hotspots. Thus, although oncogenic Ras activates genetic signaling pathways that promote invasive phenotypes, another factor should be required for the onset of the tumor invasion. Our data show that the factor is the JNK signaling, and this is consistent with previous reports showing functional cooperation of oncogenic Ras and JNK activation in cancer progression[15,39,52]. In the previous reports, *Ras^V12^* expression in combination with a mutant of *nTSG* or overactivation of JNK was used to induce invasion phenotypes in the epithelial tissues. In contrast, we showed that *Ras^V12^* expression alone can induce invasive phenotypes at a few invasive hotspots where the epithelial organization is intrinsically compromised. Our data suggest that the intrinsic mild polarity disturbance predisposes the invasion hotspot cells to activate JNK signaling by an oncogenic stimulus such as *Ras^V12^* expression and that oncogenic Ras and the JNK activation cooperatively provoke invasive behaviors of tumor cells.

Another key unanswered question is about physiological aspects of the invasion hotspots and how the spots are formed in epithelial development. The fate map of the *Drosophila* wing imaginal disc based on an elaborate implantation experiment [44] helps us to determine the developmental fate of each spot in the third instar wing discs. According to the fate map, one invasion hotspot located in the ventral hinge region is the presumptive axillary pouch (or pleural sclerite) and another one in the dorsal hinge region is the presumptive unnamed plate (or humeral plate). We still do not understand why these spots intrinsically have a disturbance of cellular arrangement patterns and a mildly compromised apicobasal organization. One plausible explanation is that those spots are mechanically distorted during morphological transformations to form a small node structure such as axillary pouch or pleural sclerite of the wing hinges. In the morphogenesis of complicated structures, developing tissues experience physical distortions to a varying degree[53]. Those mechanical distortions of cell shape, cellular membranes, cytoskeletons, or ECM, in some cases, play a key role in the control of morphogenesis, differentiation, and proliferation through mechanotransduction[54].

The invasion hotspots we identified in the wing imaginal epithelia are formed as small local spots (smaller than 10 cells in diameter) with some structural distortions. Given the conservation of epithelial cell/tissue structures in flies and mammals, it is likely that a substantial number of structurally similar spots exist in human epithelial tissues. It is also possible that tumor cells may utilize additionally occurring mutations to distort tissue structures similar to the invasion hotspots by themselves during tumor progression[11]. Future studies to identify the developmental processes by which invasion hotspots form in various types of tissues or to clarify the behaviors of different types of tumor cells in invasion hotspots will lead to a better understanding of tumor invasion mechanisms.

## 4. Materials and Methods

### 4.1. Fly Stocks and Genetics

*Drosophila* stocks were maintained by standard methods at 25°C. All fly crosses were carried out at 25°C according to standard procedures. For the generation of genetically mosaic clones, the heat-shock-activated flip-out-Gal4-UAS system[37] or mosaic analysis with a repressible cell marker (MARCM) system[36] was used. To obtain genetically mosaic mutant clones in imaginal discs, first instar larvae (48 h after egg deposition) were heat-shocked for 30-120 min at 37°C. After heat shock, larvae were maintained at 25°C until dissection of imaginal discs. To induce misexpression of *Ras^V12^* or *lgl^RNAi^* and *Ras^V12^* at the dorsal hinge region of the wing imaginal discs, *GMR17G12-Gal4* was used. The wing discs were dissected 7 days after egg deposition at 29 °C. The following fly strains were used: *scrib^1^*[55], *UAS-Ras^V12^* (BDSC #64196), *UAS-lgl-RNAi* (VDRC #51247), *UAS-Dcr-2* (BDSC #24651), *TRE-DsRed* (BDSC #59011), *UAS-bsk^DN^* (BDSC #9311), *UAS-timp* (BDSC #58708), *UAS-p35* (BDSC #5072, 5073), *GMR17G12-Gal4* (FlyLight #R17G12), *UAS-mCD8.mRFP* (BDSC #27399), *UAS-RedStinger*, *UAS-FLP*, *Ubi-p63E(FRT.STOP)Stinger* (BDSC #28281), *Vkg-GFP* (a gift from Dr. Sa Kan Yoo).

### 4.2. Detailed Genotypes for Each Experiment

**Figure 1**

(**A**−**D, F**), *hsFLP; UAS-Ras^V12^/act-Gal4, UAS-GFP; FRT82B scrib^1^*/*FRT82B tubP-Gal80*

(**G**, **I**), *hsFLP; UAS-Ras^V12^*/+*; act>CD2>Gal4, UAS-GFP*/*UAS-lgl-RNAi, UAS-dicer2*

(**H**), *hsFLP; Vkg-GFP*/*UAS-Ras^V12^; act>CD2>Gal4, UAS-RFP*/*UAS-lgl-RNAi, UAS-dicer2*

(**J**), *hsFLP;; act>CD2>Gal4, UAS-GFP*/*+*

*hsFLP; UAS-Ras^V12^*/+*; act>CD2>Gal4, UAS-GFP*/*UAS-lgl-RNAi, UAS-dicer2*

**Figure 2**

(**A**−**E**), *hsFLP; UAS-Ras^V12^*/+*; act>CD2>Gal4, UAS-GFP*/*+*

(**F**), *hsFLP;; act>CD2>Gal4, UAS-GFP*/*+*

*hsFLP; UAS-Ras^V12^*/+*; act>CD2>Gal4, UAS-GFP*/*+*

*hsFLP; UAS-Ras^V12^*/+*; act>CD2>Gal4, UAS-GFP*/*UAS-lgl-RNAi, UAS-dicer2*

**Figure 3**

(**A**−**C**), *hsFLP; UAS-Ras^V12^*/*TRE-DsRed; act>CD2>Gal4, UAS-GFP*/*UAS-lgl-RNAi, UAS-dicer2*

(**D**−**F**), *hsFLP; UAS-Ras^V12^*/*TRE-DsRed; act>CD2>Gal4, UAS-GFP*/*+*

(**G**−**H**), *hsFLP; TRE-DsRed*/+*; act>CD2>Gal4, UAS-GFP*/*+*

*hsFLP; UAS-Ras^V12^*/*TRE-DsRed; act>CD2>Gal4, UAS-GFP*/*+*

*hsFLP; UAS-Ras^V12^*/*TRE-DsRed; act>CD2>Gal4, UAS-GFP*/*UAS-lgl-RNAi, UAS-dicer2*

**Figure 4**

(**A**), *hsFLP; act>CD2>Gal4, UAS-GFP*/+*; UAS-lgl-RNAi, UAS-dicer2*/*UAS-p35*

(**B**−**C**), *hsFLP;; act>CD2>Gal4, UAS-GFP*/*+*

*hsFLP; act>CD2>Gal4, UAS-GFP*/+*; UAS-lgl-RNAi, UAS-dicer2*/*UAS-p35*

*hsFLP; UAS-Ras^V12^*/+*; act>CD2>Gal4, UAS-GFP*/*UAS-lgl-RNAi, UAS-dicer2*

(**E**−**H**), *hsFLP; UAS-p35*/*TRE-DsRed; UAS-lgl-RNAi, UAS-dicer2*/*act>CD2>Gal4, UAS-GFP*

(**I**−**L**), *hsFLP; act>CD2>Gal4, UAS-GFP*/+*; UAS-lgl-RNAi, UAS-dicer2*/*UAS-p35*

**Figure 5**

(**A**−**B**), *hsFLP; act>CD2>Gal4, UAS-GFP*/*UAS-Ras^V12^; UAS-lgl-RNAi, UAS-dicer2*/*UAS-bsk^DN^*

(**C**), *hsFLP; act>CD2>Gal4, UAS-GFP*/*UAS-Ras^V12^; UAS-lgl-RNAi, UAS-dicer2*/*UAS-timp*

(**E**), *hsFLP; UAS-Ras^V12^*/+*; act>CD2>Gal4, UAS-GFP*/*UAS-lgl-RNAi, UAS-dicer2*

*hsFLP; act>CD2>Gal4, UAS-GFP*/*UAS-Ras^V12^; UAS-lgl-RNAi, UAS-dicer2*/*UAS-timp*

**Figure 6**

(**A**), *hsFLP; UAS-Ras^V12^*/+*; act>CD2>Gal4, UAS-GFP*/*+*

(**C**), *hsFLP; UAS-Ras^V12^*/+*; act>CD2>Gal4, UAS-GFP*/*UAS-lgl-RNAi, UAS-dicer2*

(**E**), *hsFLP; UAS-Ras^V12^*/+*; act>CD2>Gal4, UAS-GFP*/*+*

*hsFLP; UAS-Ras^V12^*/+*; act>CD2>Gal4, UAS-GFP*/*UAS-lgl-RNAi, UAS-dicer2*

**Figure 7**

*w*^1118^

**Figure 8**

(**A**−**C**), *w;; GMR17G12-Gal4*/*UAS-RedStinger, UAS-FLP*, *Ubi-p63E(FRT.STOP)Stinger*

(**D**−**I**), *w; UAS-Ras^V12^*/+*; GMR17G12-Gal4, UAS-mCD8.mRFP*/+

(**J**−**L**), *w; UAS-Ras^V12^*/+*; GMR17G12-Gal4, UAS-mCD8.mRFP*/*UAS-lgl-RNAi, UAS-dicer2*

**Figure S1**

(**A**−**C**), *hsFLP; UAS-Ras^V12^*/+*; act>CD2>Gal4, UAS-GFP*/*UAS-lgl-RNAi, UAS-dicer2*

(**D**−**F**), *hsFLP;; act>CD2>Gal4, UAS-GFP*/*+*

**Figure S2**

(**A**−**C**), *hsFLP; TRE-DsRed*/+*; act>CD2>Gal4, UAS-GFP*/*+*

**Figure S3**

(**A**−**C**), *w;; GMR17G12-Gal4, UAS-mCD8.mRFP*/+

(**D**−**F**), *w; UAS-Ras^V12^*/+*; GMR17G12-Gal4, UAS-mCD8.mRFP*/*UAS-lgl-RNAi, UAS-dicer2*

### 4.3. Immunohistochemistry and Image Analysis

For analyses of genetically mosaic clones in *Drosophila* imaginal discs, larvae were chosen at the given time after clone induction, and dissected tissues were fixed in 4% formaldehyde at room temperature for 15 min. All subsequent steps for immunostaining were performed according to standard procedures for confocal microscopy[10]. The following antibodies were used: rabbit anti-cleaved *Drosophila* Dcp-1 (#9578) (1:100, Cell Signaling Technology), mouse anti-Armadillo N2 7A1 (1:40, Developmental Studies Hybridoma Bank [DSHB]), rat anti-DE-Cadherin DCAD2 (1:30, DSHB), mouse anti-MMP1 (1:1:1 mixture of 3B8, 3A6 and 5H7 were diluted 1:10, DSHB), mouse anti-*α*-Spectrin 3A9 (1:50, DSHB), mouse anti-*α*-Tubulin AA4.4 (1:100, DSHB), and rabbit anti-Laminin-γ (1:100, abcam). Alexa Fluor 488, 546, and 633 (1:400, Molecular Probes) were used for secondary antibodies. F-actin was stained by Alexa Fluor 488 and 546 Phalloidin (1:50, Molecular Probes). All samples were counterstained with DAPI (Sigma-Aldrich) for visualization of DNA. Immunofluorescence images were captured on the Olympus FV1000 or FV1200 confocal microscopes and analyzed with ImageJ. 3-D reconstructions of confocal z-stack images were rendered with ImageJ 3-D Viewer, an ImageJ plugin (B. Schmid, 2007). Signal intensities were plotted with Interactive 3-D Surface Plot, an ImageJ plugin (K.U. Barthel, 2004).

### 4.4. Quantification of pseudopodial protrusions

We defined cellular processes which project more than 5 μm from cell bodies as psudopodial protrusions. The number of psudopodial protrusions per mutant clone cluster (consisting of more than ten GFP-positive cells) were counted from five wing discs in each independent experiment for each genotype.

### 4.5. Quantification for fluorescent signal intensities

Signal intensities of TRE-DsRed or anti-MMP1 antibody staining were measured from confocal images with ImageJ. Each value of signal intensity was measured as an average intensity of a selected area of 225-square pixels (for TRE-DsRed) or 25-square pixels (excluding nuclei for MMP1) from GFP-positive clone areas and calculated as a ratio relative to control (GFP-negative areas).

### 4.6. Statistical Analysis

For data analyses, a two-tailed unpaired Student’s *t*-test was used to determine P-values. P-values less than 0.005 were considered to be significant. No statistical method was used to predetermine sample size.

## Author Contributions

Conceptualization, Y.F. and Y.T.; Methodology, Y.T.; Validation, R.K., H.T., S.D., Y.F. and Y.T.; Formal Analysis, R.K., H.T., S.D., and Y.T.; Investigation, R.K., H.T., S.D., and Y.T.; Resources, Y.F. and Y.T.; Data Curation, R.K., H.T., S.D., Y.F., and Y.T.; Writing – Original Draft Preparation, Y.T.; Writing – Review & Editing, R.K., H.T., S.D., Y.F., and Y.T.; Funding Acquisition, Y.T.

## Acknowledgments

We thank SK. Yoo, T. Igaki, the Vienna Drosophila RNAi Center, and the Bloomington Drosophila Stock Center for providing fly stocks.

## Supplementary Materials

**Figure S1**. *nTSG-Ras^V12^* double mutant cells show apical or basal extrusions in wing imaginal epithelia. (**A***−***C**) A wing disc with mosaic mutant clones coexpressing *lgl^RNAi^* and *Ras^V12^* 48 hours after clone induction. Mutant clones were marked with GFP expression (green) in (**A***−***B**). Basement membranes were labeled with anti-Laminin *γ* antibody (red) in (**A**, **C**). The images are z-stack projections of 30 confocal images of the columnar epithelial layer. Lower panels: vertical sections at a site indicated by a white line in each upper panel. A yellow arrowhead: apically extruded mutant clones. Magenta arrows: basally extruded mutant clones. PM: peripodial membrane. (**D**) A wing imaginal disc with mosaic clones expressing GFP (green) 96 hours after clone induction, stained for F-actin (red). The images are z-stack projections of 30 confocal images of the columnar epithelial layer. (**E**) F-actin detected by Phalloidin staining in the mosaic wing disc in (**D**). (**F**) Laminin detected by anti-Laminin-*γ* antibody staining (white) in the mosaic wing disc in (**D**). The images are z-stack projections of 30 confocal images of the columnar epithelial layer. Lower panels: vertical sections at a site indicated by a white line in each upper panel. Nuclei were labeled with DAPI (blue) in (**A***−***D**). Scale bars represent 50 μm.

**Figure S2.** JNK-MMP1 signaling is not endogenously activated in wing imaginal epithelia. (**A**) A wing imaginal disc with mosaic clones expressing GFP (green) 96 hours after clone induction. (**B**) JNK activation detected by TRE-DsRed in the mosaic wing disc in (**A**). (**C**) MMP1 expression detected by anti-MMP1 antibody staining (white) in the mosaic wing disc in (**A**). The images are z-stack projections of 30 confocal images of the columnar epithelial layer. White arrowheads indicate endogenous MMP1 expression in the trachea. Nuclei were labeled with DAPI (blue). Scale bars represent 50 μm.

**Figure S3.** *lgl^RNAi^-Ras^V12^* co-expression induces MMP1 upregulation in the invasion hotspot. (**A**) A wing imaginal disc with *GMR17G12-Gal4*-driven RFP expression (red) in the dorsal hinge region stained for MMP1 (green). (**B**) *GMR17G12-Gal4*-induced RFP expression pattern (red) in the wing disc (**A**). (**C**) MMP1 expression pattern (green) in the wing disc (**A**). (**D**) A wing imaginal disc with *GMR17G12-Gal4*-driven *lgl^RNAi^* and *Ras^V12^* expression in the dorsal hinge region stained for MMP1 (green). *GMR17G12-Gal4* expressing regions were labeled by RFP (red). (**E**) *GMR17G12-Gal4*-induced RFP expression pattern (red) in the wing disc (**D**). (**F**) MMP1 expression pattern (green) in the wing disc (**D**). Magenta arrowheads indicate invasion hotspots. White arrowheads indicate endogenous MMP1 expression in the trachea. Nuclei were labeled with DAPI (blue). Scale bars represent 50 μm.

