## Supplementary figures and images for "Tumor-Cell Invasion Initiates at Invasion Hotspots, an Epithelial Tissue-Intrinsic Microenvironment"

### Supplemental Figure 1

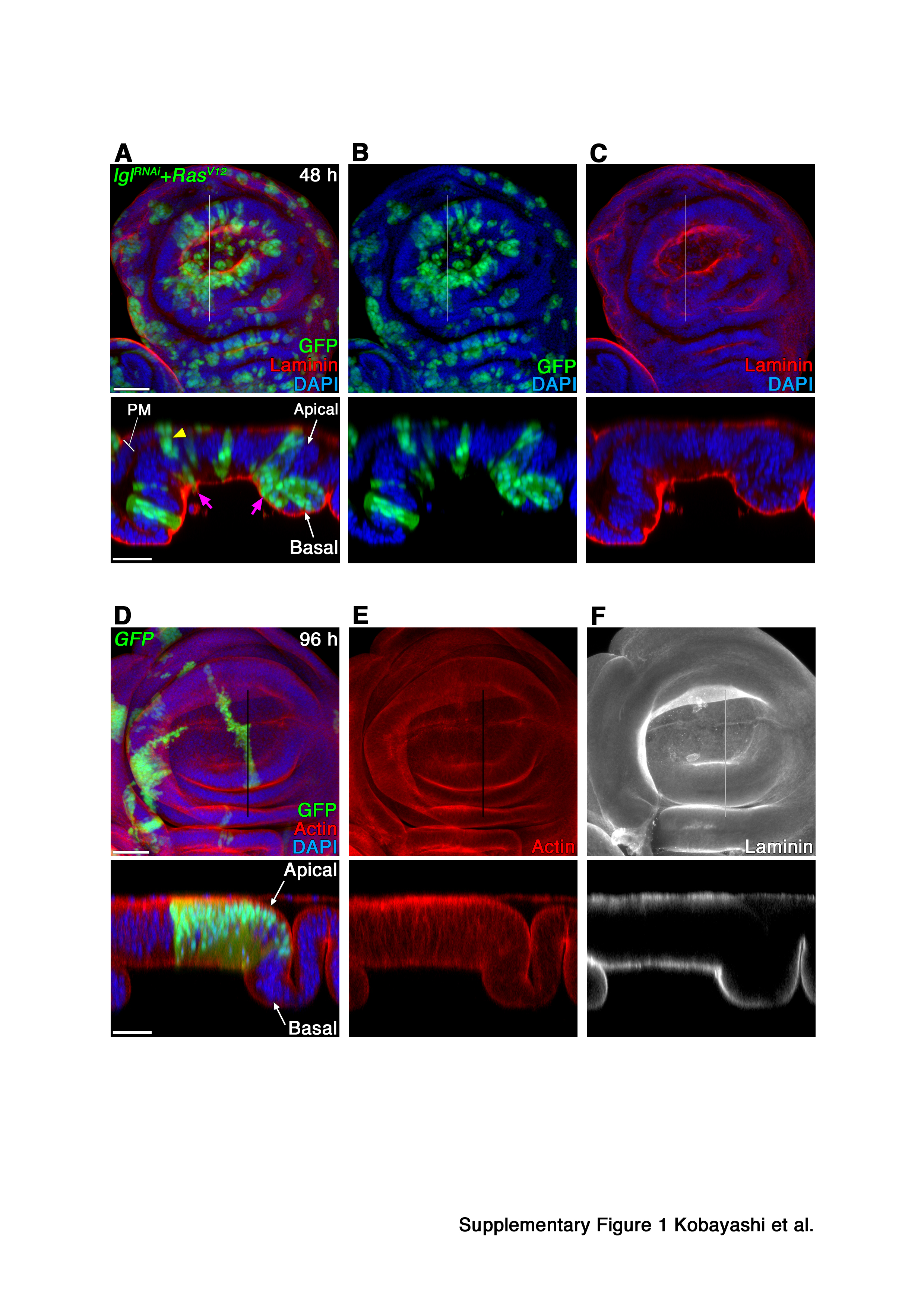

### Supplemental Figure 2

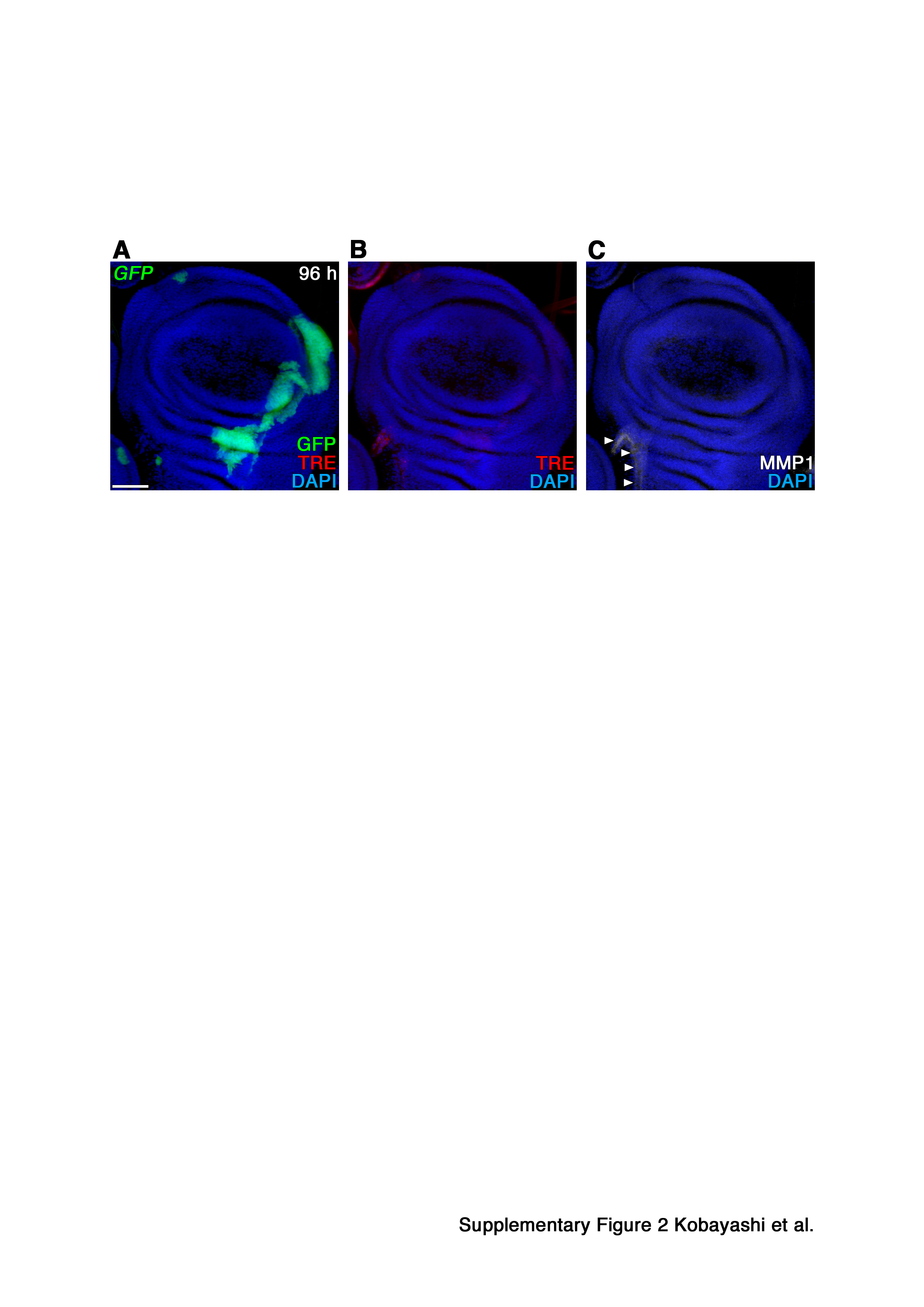

### Supplemental Figure 3

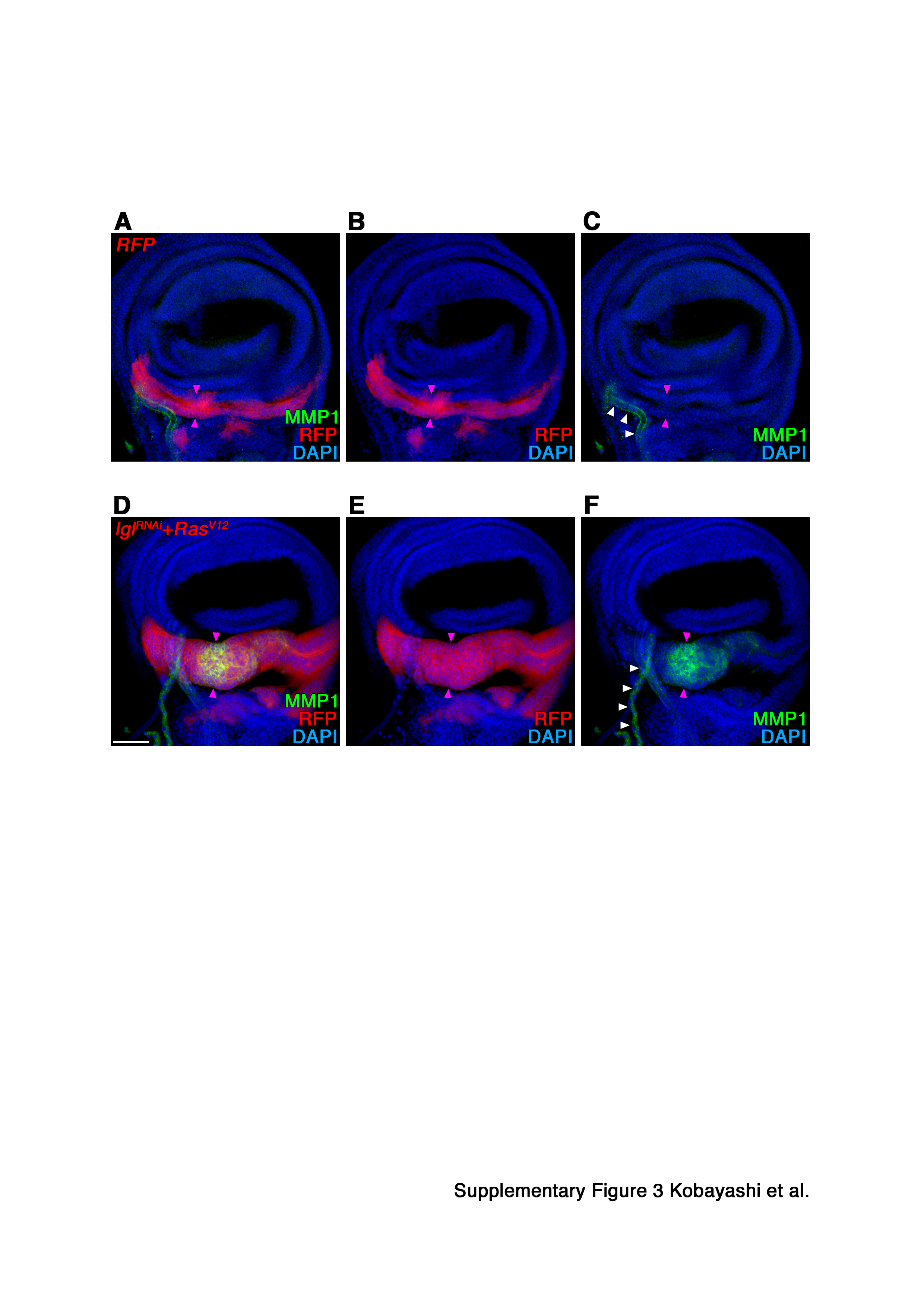
